# A genomic view of coral-associated *Prosthecochloris* and a companion sulfate-reducing bacterium

**DOI:** 10.1101/2019.12.20.883736

**Authors:** Yu-Hsiang Chen, Shan-Hua Yang, Kshitij Tandon, Chih-Ying Lu, Hsing-Ju Chen, Chao-Jen Shih, Sen-Lin Tang

**Author notes:** These authors contributed equally to this work (Yu-Hsiang Chen, Shan-Hua Yang).

## Abstract

Endolithic microbial symbionts in the coral skeleton may play a pivotal role in maintaining coral health. However, compared to aerobic microorganisms, research on the roles of endolithic anaerobic microorganisms and microbe-microbe interactions in the coral skeleton are still in their infancy. In our previous study, we showed that a group of coral-associated *Prosthecochloris* (CAP), a genus of anaerobic green sulfur bacteria, was dominant in the skeleton of the coral *Isopora palifera*. Though CAP is diverse, the 16S rRNA phylogeny presents it as a distinct clade separate from other free-living *Prosthecochloris*. In this study, we build on previous research and further characterize the genomic and metabolic traits of CAP by recovering two new near-complete CAP genomes—*Candidatus* Prosthecochloris isoporaea and *Candidatus* Prosthecochloris sp. N1—from coral *Isopora palifera* endolithic cultures. Genomic analysis revealed that these two CAP genomes have high genomic similarities compared with other *Prosthecochloris* and harbor several CAP-unique genes. Interestingly, different CAP species harbor various pigment synthesis and sulfur metabolism genes, indicating that individual CAPs can adapt to a diversity of coral microenvironments. A novel near-complete SRB genome—*Candidatus* Halodesulfovibrio lyudaonia—was also recovered from the same culture. The fact that CAP and various sulfate-reducing bacteria (SRB) co-exist in coral endolithic cultures and coral skeleton highlights the importance of SRB in the coral endolithic community. Based on functional genomic analysis of *Ca*. P. sp. N1 and *Ca*. H. lyudaonia, we also propose a syntrophic relationship between the SRB and CAP in the coral skeleton.

**Importance:** Little is known about the ecological roles of endolithic microbes in the coral skeleton; one potential role is as a nutrient source for their coral hosts. Here, we identified a close ecological relationship between CAP and SRB. Recovering novel near-complete CAP and SRB genomes from endolithic cultures in this study enabled us to understand the genomic and metabolic features of anaerobic endolithic bacteria in coral skeletons. These results demonstrate that CAP members with similar functions in carbon, sulfur, and nitrogen metabolisms harbor different light-harvesting components, suggesting that CAP in the skeleton adapts to niches with different light intensities. Our study highlights the potential ecological roles of CAP and SRB in coral skeletons and paves the way for future investigations into how coral endolithic communities will respond to environmental changes.

## Introduction

Microbial symbionts in reef-building corals, which support a variety of marine life, reside in the mucus, tissue, and skeleton of diverse corals, influencing health of its host coral (1, 2). Microbial symbionts comprise bacteria, archaea, algae, fungi, and viruses, and their composition is influenced by their host corals’ genetic factors and dynamic environmental conditions (3). They can help corals prevent or mitigate diseases and benefit corals by involving them in carbon, nitrogen, and sulfur cycles (4). For example, coral dominant dinoflagellate *Symbiodinium* can fix carbon dioxide and provide corals with organic compounds (5). On the other hand, *Cyanobacteria* can fix nitrogen and provide the coral *Montastraea cavernosa* with a nitrogen source (6).

Compared to aerobic microorganisms, the role of anaerobic microorganisms in coral is not well understood. Previous studies found green sulfur bacteria (GSB) in a wide range of corals, including *Porites lutea, Platygyra carnosa, Montastraea faveolata* and *Montipora venosa* (7-10). In addition, our previous study found that *Prosthecochloris*, a GSB genus, was dominant in skeletons of the coral *Isopora palifera*, forming a distinct green color region beneath the coral tissue (11), although the algae *Osterobium* were previously thought to be the main microbial contributor to coral green layers (11-13). Moreover, nutrients generated from microorganisms in the coral skeleton were shown to be potential alternative sources of energy and nutrients (14, 15). Therefore, the *Prosthecochloris* dominant in green layers may also be associated with stony coral health.

Most GSB are obligate anaerobic photoautotrophic bacteria that use the reverse tricarboxylic acid (rTCA) cycle to fix carbon dioxide (16). During photosynthesis, the majority of them utilize reduced sulfur compounds as electron donors, while some— including *Chlorobium ferrooxidans* and *C. phaeoferrooxidans*—use ferrous iron (17-19). Furthermore, some GSB are capable of obtaining reduced sulfur compounds through a syntrophic interaction with sulfur-reducing bacteria (SRB), such as *Desulfuromonas acetoxidans* (20). On the other hand, many GSB can fix nitrogen gas, which they use for growth (16). GSB are found in various anoxic environments—including freshwater, hot springs, and seawater—and some of them are adapted to light-limited environments (16). Among GSB, *Prosthecochloris* is mainly present in marine environments and has the ability to tolerate high salinity (16).

Though *Prosthecochloris* and most other GSB have been isolated as free-living bacteria (16), our previous study used amplicon and whole-metagenome analyses and found that *Prosthecochloris* is dominant in green layers of coral *Isopora palifera* skeletons, suggesting that the bacteria can interact with eukaryotic hosts and various bacteria (11, 13). Through a phylogenetic analysis of the 16S rRNA gene, we found that, although *Prosthecochloris* from coral were diverse, they could be classified into a monophyletic clade separate from other free-living *Prosthecochloris*. Hence, we proposed a group of coral-associated *Prosthecochloris* (CAP) (11). Furthermore, based on a gene-centric metagenome analysis, we proposed that CAP can fix nitrogen and nutrient cycling occurs in the coral skeleton.

The role of endolithic microbiomes in the coral reef system has been overlooked (21). To provide detailed insights into the ecological roles of CAP and microbe-microbe interactions in the coral skeleton, high-quality genomes of endolithic microbes are needed. The genome for the CAP *Candidatus* Prosthecochloris A305, which we identified by metagenome-binning, is only 79% complete. Other metagenomic bins identified were highly contaminated with other species. These results hindered our understanding of the metabolic features of CAP and illuminated syntrophic relationships between CAP and other microorganisms in the coral skeleton. Using an anaerobic culture approach, three endolithic cultures dominated by CAP were successfully obtained. The cultures, containing purer and more simplified communities and sufficient genomic DNA, enabled us to obtain the high-quality genomes of CAP and other companion bacteria using whole-metagenome sequencing approach. In this study, we recovered two near-complete CAP genomes from the metagenomes of the coral endolithic cultures. These new genomes allowed us to compare functional genomic and phylogenetic features in CAP and to elucidate its diversity. Moreover, we also identified a novel, predominant sulfate-reducing bacteria (SRB) genome from the same cultures. Based on functional genomic analysis in these genomes, we propose a syntrophic relationship between CAP and SRB in the coral skeleton.

## Results

### High-quality bins recovered from coral endolithic cultures

Reads from coral endolithic cultures (N1, N2, and N3) were individually *de novo* assembled and binned, yielding 5, 5, and 4 bins, respectively (Table 1). Bins from cultures had similar taxonomic profiles, dominated by *Prosthecochloris*-related bins in N2 and N3 and *Ilyobacter*-related bins in N1 (Table S1). On the other hand, *Halodesulfovibrio*-related bins were the most abundance sulfate-reducing bacterial bins in the three coral endolithic cultures. Other genera represented in bins were *Marinifilum, Pseudovibrio*, and *Desulfuromonas*, which were present in two of the three cultures. Among the total 14 bins identified, nine were high-quality (>90% complete and <5% contamination). The *Prosthecochloris*-related bins had particularly high quality (>98.8% complete) and low contamination (<1.5%); *Halodesulfovibrio*-related bins in N3 was also near-complete (99.41%) with very low contamination (0.26%) (Table 1). Both *Prosthecochloris-* and *Halodesulfovibrio-*related bins lacked strain heterogeneity.

**Table 1.** Qualities and putative taxon of each bins in metagenome from N1, N2, and N3 cultures.

| Bin Id | putative taxonomy | Completeness (%) | Contamination (%) | Strain heterogeneity | Genome size (bp) | # contigs | N50 | Mean contig length (bp) | Longest contig (bp) | GC | # predicted genes |
| --- | --- | --- | --- | --- | --- | --- | --- | --- | --- | --- | --- |
| n1_1 | *Marinifilum fragile* | 99.46 | 2.15 | 0 | 4,632,452 | 67 | 126,188 | 69,141 | 306,114 | 35.7 | 3,843 |
| n1_2 | *Desulfuromonas* sp. | 91.15 | 2.58 | 0 | 4,431,711 | 306 | 19,053 | 14,482 | 125,708 | 55.0 | 4,146 |
| n1_3 | *Halodesulfovibrio* sp. | 100 | 0.56 | 33.33 | 4,215,690 | 43 | 163,138 | 98,039 | 320,254 | 45.1 | 3,684 |
| n1_4 | *Ilyobacter* sp. | 94.38 | 1.12 | 0 | 2,867,017 | 124 | 32,721 | 23,121 | 160,652 | 36.3 | 2,715 |
| n1_5 | *Prosthecochloris marina* | 99.43 | 1.37 | 0 | 2,785,587 | 24 | 205,628 | 116,066 | 495,280 | 47.0 | 2,648 |
| n2_1 | *Halodesulfovibrio* sp. | 97.93 | 2.73 | 86.67 | 3,681,226 | 182 | 29,684 | 20,226 | 89,738 | 45.1 | 3,294 |
| n2_2 | *Desulfuromonas* sp. | 63.38 | 2.58 | 40 | 2,938,736 | 622 | 4,902 | 4,724 | 22,680 | 55.8 | 3,037 |
| n2_3 | *Ilyobacter* sp. | 96.63 | 1.12 | 0 | 2,896,854 | 127 | 33,041 | 22,809 | 160,613 | 36.3 | 2,742 |
| n2_4 | *Prosthecochloris* sp. | 99.45 | 0.82 | 0 | 2,627,088 | 52 | 65,875 | 51,404 | 309,532 | 47.4 | 2,545 |
| n2_5 | *Desulfovibrio bizertensis* | 80.85 | 1.18 | 0 | 2,284,992 | 440 | 5,709 | 5,193 | 28,696 | 52.6 | 2,379 |
| n3_1 | *Marinifilum* sp. | 99.19 | 2.42 | 0 | 5,498,267 | 61 | 142,436 | 90,135 | 543,023 | 35.9 | 4,546 |
| n3_2 | *Pseudovibrio* sp. | 85.04 | 0.79 | 0 | 5,165,768 | 718 | 8,788 | 7,194 | 35,672 | 50.0 | 5,091 |
| n3_3 | *Halodesulfovibrio* sp. | 99.41 | 0.26 | 0 | 3,714,212 | 81 | 77,081 | 45,854 | 159,330 | 44.9 | 3,295 |
| n3_4 | *Prosthecochloris* sp. | 98.90 | 0.82 | 0 | 2,630,645 | 50 | 79,255 | 52,612 | 225,631 | 47.4 | 2,545 |

### Novel near-complete coral-associated *Prosthecochloris* (CAP) draft genomes from coral endolithic cultures

The results of the GTDB-Tk taxonomy assignment showed that all bins were closest to *Prosthecochloris marina* V1. Interestingly, *Prosthecochloris*-related bins in N2 and N3 shared only 90% Average Nucleotide Identity (ANI) with *Prosthecochloris* marina V1 (Fig. S1), which is below the 95% ANI cutoff, a frequently used standard for species delineation (22). On the other hand, the ANI between *Prosthecochloris*-related bins in N2 and N3 was 99.9%, suggesting that the bins were identical, and these bins were named *Candidatus* Prosthecochloris isoporaea. The draft genome of *Ca*. P. isoporaea was 2.6 Mb with 47.4% GC, which is within the range of *Prosthecochloris* genomes (2.4 - 2.7 Mb with 47.0 - 56.0% GC). The N50 of the draft genomes in N2 and N3 were 65 and 92 kbp, respectively. The contig numbers were 51 and 49, and the longest contig was 309 and 225 kbp, respectively. The longest contig of the *Ca*. P isoporaea genome in N2 was larger, so this genome was used as the representative genome for all downstream analysis.

The ANI between the *Prosthecochloris*-related bin in N1 and *Prosthecochloris marina* V1 was 99%, suggesting that these genomes belong to the same species. The bin was named *Candidatus* Prosthecochloris sp. N1. Its genome size was 2.7 Mb, with 23 contigs and a 47.0% GC ratio, which is consistent with the genome of *Prosthecochloris* marina V1 (23).

The ANI between these newly identified genomes and other *Chlorobiaceae* members was also determined (Fig. S1). *Ca*. P. isoporaea and *Ca*. P. sp. N1 shared the highest ANI value with *Candidatus* Prosthecochloris sp. A305 (∼79%) and *Candidatus* Prosthecochloris korallensis (∼80%), which were both previously identified from the coral metagenomes and defined as part of the coral-associated *Prosthecochloris* (CAP) group (11). Furthermore, the genomes of *Candidatus* Prosthecochloris sp. A305 and *Candidatus* P. korallensis were most similar (82% ANI) (Fig. S1). These results indicated high genomic similarities between the members of CAP. The other *Chlorobiaceae* closest to CAP were *Prosthecochloris* sp. GSB1 and *Chlorobium phaeobacteroides* BS1, later annotated as *Prosthecochloris phaeobacteroides* BS1 (7).

### Phylogenetic tree of CAP and other green sulfur bacteria

To determine the phylogenetic relationship between CAP and other members of *Chlorobiaceae*, 16S rRNA gene sequences of CAP-related genomes and other *Chlorobiaceae* were used to reconstruct phylogenetic trees (Fig. 1A). The analysis also included *Prosthecochloris*-related Operational Taxonomic Units (OTU) (at species-like level), which we identified from the green layer of coral *Isopora palifera* (11); bin-3, which was recovered from metagenomes in the green layer of *Isopora palifera* (11); and one uncultured clone isolated from the coral *Montastraea faveolata* (24). All CAP members were grouped into the same clade, and the clade closest to it contained other free-living *Prosthecochloris*. The tree based on FMO, a unique photosynthetic-related protein in *Chlorobiaceae*, also classified the CAP members into the same clade, with the addition of *Chlorobium phaeobacteroides* BS1 and *Prosthecochloris* sp. GSB1 (Fig. 1B). In addition, to more confidently establish the evolutionary relationships, we also used concatenated protein sequence alignments of common single-copy genes in these genomes to construct the tree. The results demonstrated that the CAP forms a unique clade, irrespective of the sequences used (Fig. 1C). These congruent results indicate that CAP have a unique evolutionary origin.

**FIG 1.**
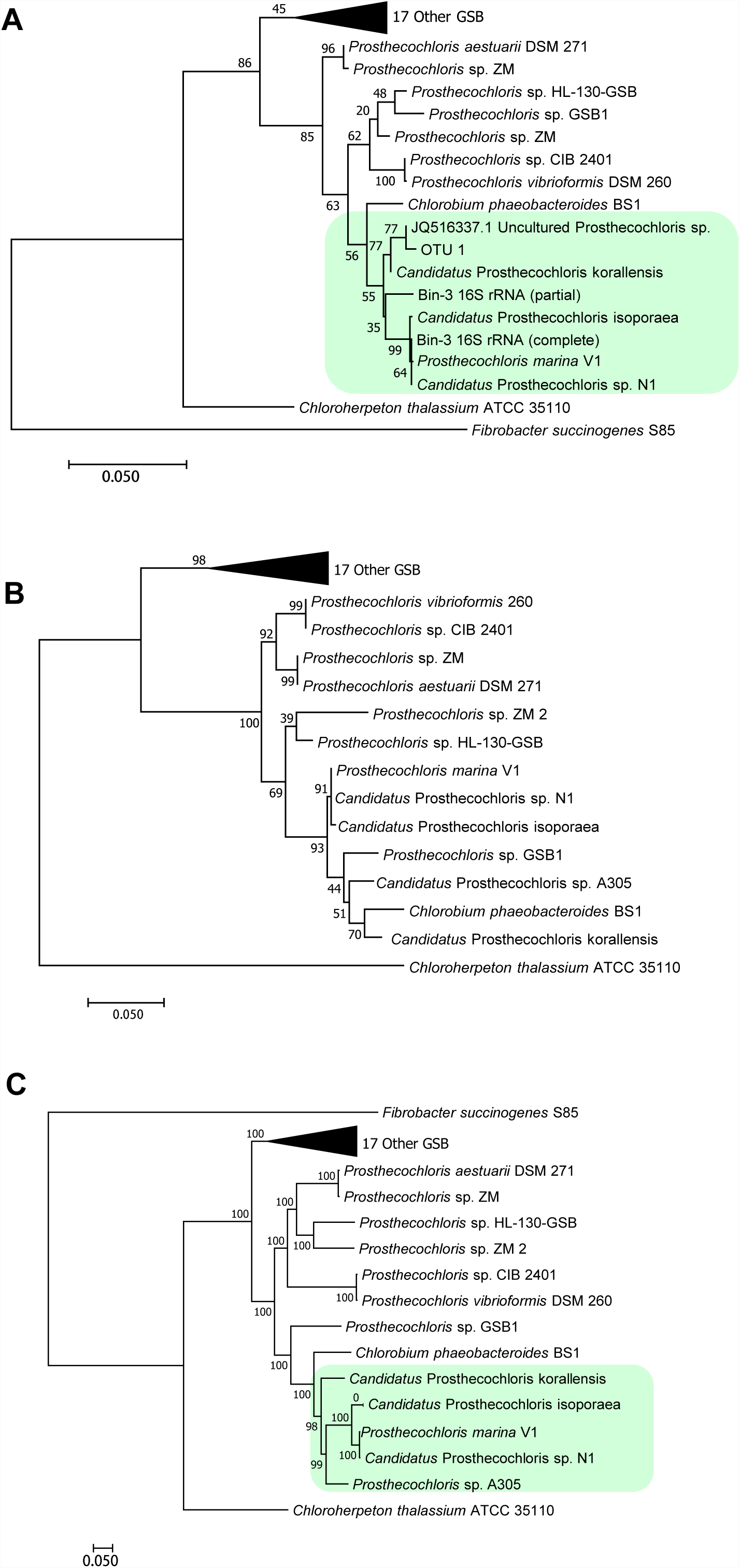
Molecular phylogenetic analysis of green sulfur bacteria. The phylogenetic trees of 16S rRNA (A), FMO protein (B), and protein sequences of concatenated single-copy genes (C) were constructed by the maximum-likelihood method with 1000 bootstraps. 27 green sulfur bacteria genomes in the RefSeq database and coral-associated GSB genomes were used to construct the tree. Other GSB included 12 *Chlorobium*, 1 *Pelodictyon*, and 4 *Chlorobaculum*. The genome and 16S rRNA sequences of *Fibrobacter succinogenes* S85 were used as the outgroup.

### Pan-genome analysis of *Prosthecochloris*

Pan-genome analysis was conducted to understand the core-accessory relationships in the genus *Prosthecochloris*. The plot of pan-genome size along the number of genomes indicated that the pan-genome is open (Fig. S2A). The *Prosthecochloris* genomes share 442 core genes (Fig. S2B). The number of genes absent only in *Candidatus* Prosthecochloris sp. A305 is 122, which may indicate that the draft genome is incomplete. The COG and KEGG classification of the core, accessory, and unique proteins revealed that the translation, energy production, and amino acid metabolism categories had higher proportions of core proteins than accessory or unique proteins (Fig. S3A and B). On the other hand, the drug resistance, secondary metabolite biosynthesis, DNA replication, and membrane transport categories had higher proportions of accessory and unique proteins (Fig. S3A and B). The phylogeny of concatenated alignment of core protein sequences grouped CAP members in the same clade (Fig. S4), with *P*. sp GSB1 and *C. phaeobacteroides* BS1 as closest relatives. The CAP clade contained 213 clade-specific accessory genes. In addition, we also found 80 genes present in all CAP genomes, except that of A305. The 213 accessory genes and these 80 genes were searched using BLASTn against the NCBI RefSeq database. The results showed that, although most genes had orthologue genes in other *Chlorobiaceae* members, some were unique to CAP members (Table S2). It is noteworthy that the putative gene sources of many BLASTn top hits were from sulfate-reducing bacteria. Moreover, the *dN/dS* ratio of these genes were < 0.3, indicating that the changes in amino acid sequences in these gene coding sequences were deleterious.

### Metabolic characteristics of CAP

The KEGG annotation by BlastKoala revealed that all the CAP members have nitrogen fixation genes—except for *Ca*. P. A305—and lack the genes for dissimilatory nitrate reduction pathway and denitrification—except for *Ca*. P. korallensis, which contains genes responsible for converting nitrite to ammonia (Table S3). For the carbon metabolism pathway, all the CAP members have a complete gene repertoire for the rTCA cycle—except for *Ca*. P. A305, which lacks the *idh* gene. On the other hand, the gene encoding phosphoenolpyruvate carboxylase (*ppc*) is only present in *Ca*. P. A305 and *Ca*. P. korallensis and the carbon monoxide dehydrogenase coding gene (*cooF* or *cooS*) is only present in *Ca*. P. korallensis and *Ca*. P. sp. N1.

For the sulfur metabolism pathways, *sqr* and *fccAB*—encoding sulfide-quinone reductase and sulfide dehydrogenase, respectively—were identified in all CAP members. Complete dissimilatory sulfate reduction (DSR) and thiosulfate reductase pathway encoding genes were identified in all members of CAP except *Ca*. P. A305. In addition, the genomes of *Ca*. P. isoporaea and *Ca*. P. sp. N1 also contained all genes in the assimilatory sulfate reduction and thiosulfate-oxidizing Sox enzyme systems, except for the *soxCD* genes.

Distinct colors of the N1 (green) and N2 (brown) cultures led us to hypothesize that CAP can harbor different bacteriochlorophylls (BChl), as a previous study showed that brown-color GSB have BChl *e* (19). The KEGG results showed that all CAP members have the genes to synthesize BChl *a*, BChl *b*, and BChl *d* from chlorophyllide *a* (Table S3), but the *bciD* gene—encoding the enzyme that converts bacteriochlorophyllide *c* to bacteriochlorophyllide *e*—is only present in *Ca*. P. isoporaea. Moreover, our previous analysis of the absorption spectrum revealed the presence of BChl *e* in the N2 culture only (11). These results implied that the presence of *bciD* gene might enable *Ca*. P. isoporaea to synthesize BChl *e*, suggesting that the differences in genes responsible for pigment synthesis could be responsible for the color difference in the N1 and N2 cultures.

The transporter systems in CAP were also identified by BlastKoala (Table S3). The results demonstrate that CAP have the ABC transporter systems for transporting molybdate, nucleoside, phospholipid, phosphate, lipoprotein, lipopolysaccharide, and cobalt. In addition, sulfate, ammonium, and drug/metabolite transporters were also identified by annotation in transportDB 2.0 (DATA SET S1).

### Recovered novel sulfate-reducing bacteria genome in coral endolithic cultures

Our binning results showed that the *Halodesulfovibrio*-related bin was present in all coral endolithic cultures, and the bin in N3 is nearly complete (99.41%) and has very low contamination (0.26%) (Table 1). The closest available genome to this bin is *Halodesulfovibrio marinisediminis*, with an ANI of 84.1%, suggesting that the bin belongs to a novel species. Hence, the bin was renamed as *Candidatus* Halodesulfovibrio lyudaonia. The total length of the draft genome is 3.7 Mb, comprising 81 contigs with a 44.9% GC ratio.

The ANI between the genomes of existing *Halodesulfovibrio* species and *Ca*. H. lyudaonia was 83 to 84%. As *Halodesulfovibrio* originally belonged to the *Desulfovibrio* genus, the ANI between *Desulfovibrio* and *Ca*. H. lyudaonia was also determined, which demonstrated that *Ca*. H. lyudaonia and some *Desulfovibrio* species share >70% ANI. The phylogenetic analysis of 16S rRNA and whole-genome similarity revealed that the *Halodesulfovibrio* could be separated from *Desulfovibrio* as a monophyletic clade (Fig. S5A and B). Besides, the 16S rRNA analysis also showed that *Ca*. H. lyudaonia and *Halodesulfovibrio*-related 16S rRNA in the N1 culture could be classified into a clade with *H. marinisediminis* and *H. spirochaetisodalis* (Fig. S5A).

The genomic analysis within sulfur metabolism revealed that all the existing *Halodesulfovibrio* and *Ca*. H. lyudaonia have dissimilatory sulfate reduction and *sqr* genes (Table S4). For the nitrogen metabolism, the nitrogen-fixation genes were only identified in *H. aestuarii*, and denitrification and nitrate reduction-related genes were absent in all genomes (Table S4). For carbon metabolism, genes participating in glycolysis and ethanol fermentation were present in all *Halodesulfovibrio*. Moreover, all genomes contained multiple genes encoding formate dehydrogenase, which helps convert formate to CO_2_.

The transporter gene analysis revealed the existence of molybdate, nucleoside, phospholipid, phosphate lipopolysaccharide, cobalt, phosphonate, glutamine, branched-amino, zinc, and tungstate transporter genes in *Halodesulfovibrio* (Table S4). Furthermore, the general L-amino acid and sulfate transporter genes were also identified in the *Ca*. H. lyudaonia. Different *Halodesulfovibrio* species contained various secretion systems. *Halodesulfovibrio* have genes responsible for the Type II secretion system, twin-arginine translocation pathway, and general secretory pathway (Table S4). Apart from these systems, the *Ca*. H. lyudaonia also had genes involved in the Types III and VI secretion systems.

## Discussion

In this study, we used genomic and functional genomics analyses to characterize coral-associated *Prosthecochloris* (CAP) and a companion sulfate-reducing bacterium. Two near-complete and high-quality CAP draft genomes were recovered from coral endolithic cultures, including one novel species. The genomic and functional analysis of existing CAP members revealed a functional diversity between the members, in spite of their phylogenetic closeness and genome similarities. Along with CAP, sulfate-reducing bacteria (SRB) were also common in endolithic cultures, indicating a potential symbiotic relationship between the groups. Hence, a near-complete draft genome of a novel species in *Halodesulfovibrio*—a common SRB genus in coral endolithic cultures—was also recovered and functional genomics analysis performed. Based on the metabolic features of the CAP and SRB genomes, a putative syntrophic interaction between the *Halodesulfovibrio* and CAP was proposed.

### CAP formed a monophyletic clade and shared several CAP-specific genes

*Prosthecochloris* is the only green sulfur bacterial genus found in green layers of coral skeleton to date. Furthermore, CAP can be phylogenetically separated from other free-living *Prosthecochloris*, suggesting that they share certain common features enabling them to live in diverse microenvironments of the coral skeleton. Interestingly, pan-genome analysis identified several genes that were unique to CAP. The similarity search results revealed that most of these genes were from SRB, suggesting a close ecological relationship between SRB and CAP members and maybe even a history of horizontal gene transfer. These CAP-unique genes had a low ratio of nonsynonymous to synonymous substitutions (*dn/ds*<1), indicating that these genes underwent purifying selection; therefore, mean the changes in the overall amino acid sequences of these genes would decrease bacteria fitness.

We propose two hypotheses about the ancestor of CAP. First, it acquired these genes while living in coral skeletons, and these genes were selected for. Second, it lived in other microbial communities and, after acquiring the above mentioned genes, gained fitness to live in coral environments. For example, among the CAP-specific genes, we found that there is a tripartite ATP-independent periplasmic transporter (TRAP transporter) gene cassette that includes permease and a substrate-binding subunit. TRAP is a protein family involved the bidirectional transport of a wide range of organic acids (25). CAP could potentially use this transport system to acquire important nutrients from the specific coral-built environment.

### CAP possess different photosynthetic machinery

Green sulfur bacteria (GSB) are obligate anaerobic photoautotrophs that use light as an energy source to grow (19). Photosynthesis occurs in self-assembly light-harvesting complexes called chlorosomes, which comprise different types of bacteriochlorophyll (BChl) pigments (19). Though all GSB have BChl in their reaction centers, different members have different antenna pigments, resulting in different colors (16). The major BChls in GSB, including BChl *c, d*, or *e*, have different absorption peaks. Green-colored GSB have BChl *c* or *d*, and brown-colored GSB contain BChl *e* in the chlorosome (16). The brown-colored GSB were shown to be well adapted to light-limited environments, such as deeper waters (19). Moreover, a previous study revealed that light conditions in a lake may determine which color of GSB will is the dominant group (16, 26).

The coral endolithic cultures N1 and N2, dominated by CAP, were green- and brown-colored, respectively. Our previous study confirmed the presence of BChl *c* and lack of BChl *e* peak in the N1 culture, from which *Candidatus* P. sp. N1 was recovered (11). On the other hand, the BChl *e* was present in the N2 culture, from which *Candidatus* P. isoporaea was identified. The functional genomics analysis in this study suggests that the lack of the *bciD* gene, which participates in BChl *e* biosynthesis, may account for the absence of BChl *e* in *Ca*. P. sp. N1, leading to the green coloration (27). This result suggests that CAP members may possess different photosynthetic machinery, which can help which species is dominant under different light conditions in coral skeleton microenvironments.

Multiple factors contribute to the variation in light availability of a skeleton microenvironment, including individual differences in skeleton pore size and skeleton structures owing to genetic differences or dynamic environmental factors (28). Light availability also varies at the different depths of the coral tissue (29). Hence, we hypothesize that the individual difference in skeleton structures and the depth of microhabitat in coral skeleton will influence the distribution of different CAP species. For instance, deeper sections of the skeleton with less light could be dominated by brown-colored CAP, while the regions closer to the surface of coral tissue may be dominated by green-colored CAP (Fig. 2). Confirming this hypothesis requires further investigating pigment contents by determining absorbance spectra in the different sections of a single coral skeleton to establish whether there is any correlation between the distribution of the two specific groups and the depth of the skeleton region.

**FIG 2.**
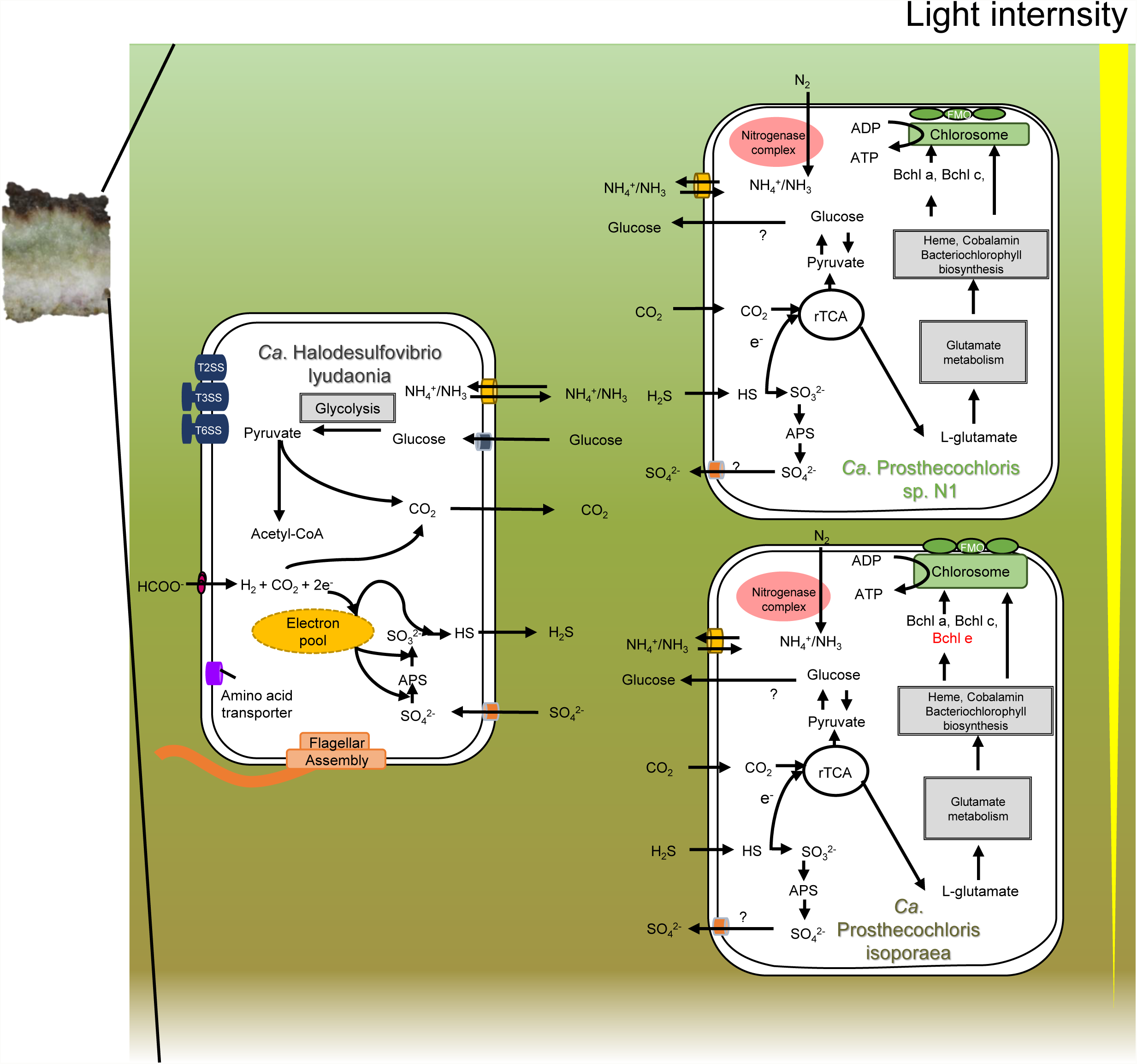
Putative syntrophic interaction between CAP and *Ca*. H. lyudaonia. Brown-colored *Ca*. P. isopraea dominated the lower section of the coral skeleton while green-colored *Ca*. P. sp. N1 dominated the upper lower section. The light intensity decreased with depth into the skeleton. The exchange of carbon, sulfur, and nitrogen compounds are denoted; important transports are indicated based on the genome annotation. The detailed model is described in the discussion.

### Sulfur metabolism in CAP

Most GSB species obtain electrons by oxidizing sulfide, sulfur, and thiosulfate for carbon fixation (30, 31). Among oxidative sulfur metabolism pathways, the Sox enzyme system—by which bacteria oxidize thiosulfate—is common. However, using thiosulfate as an electron donor and Sox gene clusters are only found in some GSB (32). In addition, GSB do not have the SoxCD complex, a part of the Sox system that is integral for oxidizing thiosulfate to sulfate in many other bacteria; instead, the function of SoxCD is replaced by the dissimilatory sulfate reduction (DSR) system in GSB (16, 33, 34). Moreover, many GSB use the DSR system to oxidize polysulfide to sulfite. Thus, in GSB, the DSR system is required to complete the oxidation of sulfur compounds. In CAP, *Ca*. P. isoporaea and *Ca*. P. sp. N1, identified from the coral skeleton, contain all genes involved in DSR and the Sox system—except for *soxCD*—indicating that GSB can obtain electrons by oxidizing sulfide, sulfite, and thiosulfate, which is similar to the way that *Chl. tepidum* operates (35). However, *Ca*. P. korallensis, identified from homogenized corals, only have the DSR system. With the DSR system, GSB are better able to utilize reduced sulfur compounds, which might confer additional advantages in sulfide- and energy-limited conditions. However, *Ca*. P. korallensis lacks the Sox system. This may due to the differences in the availability of sulfur compounds inside corals, which contribute to the diverse sulfur metabolism in CAP or the incompleteness of *Ca*. P. korallensis genome.

In some anaerobic systems, the syntrophic interaction between GSB and sulfur-reducing bacteria (SRB) occurs because sulfate produced by GSB is used as an electron acceptor in SRB, and biogenic sulfide produced by SRB is used as an electron donor in GSB (20). The binning results and our previous 16S rRNA gene-based analysis in endolithic cultures revealed the presence of potential SRB including *Halodesulfovibrio, Desulfovibrio*, and *Desulfuromonas*. These bacteria are common in the skeleton of *Isopora palifera* (11). In the three endolithic cultures, the SRB was predominant in metagenomic sequencing, suggesting that it 1) is the main group providing reduced sulfur compounds as electron donors for CAP in cultures and 2) plays the synergetic role in the endolithic community in coral skeletons.

### A novel sulfate-reducing bacterium genome identified from coral endolithic cultures

Our metagenome analyses demonstrated the relationship between CAP and SRB. The most abundant SRB in our coral endolithic cultures is *Halodesulfovibrio*, which is present in all cultures and also in green layers. Here, we recovered a high-quality near-complete draft genome of a novel species *Candidatus* Halodesulfovibrio lyudaonia. *Halodesulfovibrio* was classified as a novel genus separated from *Desulfovibrio* according to the differences in genome, phylogeny, and phenotype in 2017 (36-38). There are currently only four available species and genomes, which were all identified from marine habitats, including sediment and oxygen minimum zone water columns. Ours is the first study to find that *Halodesulfovibrio* might have a relationship with its eukaryotic host and may have syntrophic relationship with other bacteria.

Previous studies revealed that *Halodesulfovibrio* can use sulfate or sulfite as electron acceptors (38). The presence of all genes involved in the DSR system indicates that these bacteria use this pathway to reduce sulfur compounds (Table S4). In addition, some SRB can also fix nitrogen, such as *Firmicutes* and *Deltaproteobacteria* (39). In our analysis, nitrogen fixation genes were absent in all *Halodesulfovibrio* except *H. aestuarii* (Table S4). However, we also found that bacteria containing the gene encoding L-amino acid and ammonia transporters can be used to obtain organic nitrogen.

### Putative syntrophic interaction between diverse CAP and *Halodesulfovibrio*

Previously, we proposed a general syntrophic interaction based on gene-centric approach with metagenomes of coral skeleton (11). Here, using several high-quality and near-complete draft genomes from endolithic cultures, we identified CAP and SRB species that participate in this syntrophic interaction. Moreover, the high-quality draft genomes also allowed us to characterize communities and interactions in a more accurate and detailed manner. The recovered genomes highlight the diversity in CAP and the complex interactions in the community (Fig. 2).

Brown-colored CAP can adapt to low-light microenvironments, and therefore may dominate deeper sections of the skeleton, while green-colored CAP may dominate the sections closer to the coral tissue, which are exposed to relatively higher light intensity. On the other hand, the presence of *Halodesulfovibrio* in all endolithic cultures—along with both brown- and green-colored CAP—suggests that *Halodesulfovibrio* may be distributed across different sections and interact with both colors of CAP. We suggest that both CAP species occupy their niches via diversified pigment compositions, and both interact in a syntrophic manner with *Halodesulfovibrio*.

During photosynthesis, these CAP obtain CO_2_ released by *Halodesulfovibrio* and other heterotrophs. To fix carbon through the rTCA cycle, CAP obtains sulfide from *Halodesulfovibrio* as an electron donor, while the *Halodesulfovibrio* obtain oxidized sulfur compounds released from CAP and reduce them using electrons from the conversion of formate to CO_2_. Therefore, CAP and *Halodesulfovibrio* provide each other with sulfur resources in the coral skeleton.

Being the most dominant nitrogen fixers, CAP fixes dinitrogen into ammonium, which can be bi-directionally diffused across the cell membrane into the microenvironment by the ammonium transporter. Although genes involved in nitrogen fixation are absent in *Halodesulfovibrio*, they can take up ammonium through an ammonium transporter, which might serve as a potential nitrogen source. Hence, we suggest that CAP plays an essential role in nitrogen fixation in the community.

## Conclusion

Though the skeleton microbiome may contain nutritional sources and facilitate the recovery of unhealthy coral (15), its importance in the coral skeleton has been overlooked, and the interactions inside the community are poorly studied due to methodological limitations (21). Here, our genomic analysis of endolithic cultures helps us better characterize the community and investigate the interaction between coral and the endolithic microbiome.

Endolithic cultures provide several near-complete and precise genomes to study endolithic communities. Genomic analysis revealed that members of CAP share a common origin and contain several CAP-specific genes, indicating that certain differences exist between CAP and other free-living *Prosthecochloris*. These differences imply that coral and CAP have a symbiotic relationship, but future investigations into metabolic exchanges between CAP and the coral host are needed to confirm this. On the other hand, functional genomic analysis revealed the diversity of pigments synthesized in CAP, suggesting that 1) individual members of CAP adapt to different microenvironments in the skeleton and 2) there is spatial heterogeneity in the microbiome. Along with CAP, the predominance of *Halodesulfovibrio* indicates that it is ecologically important in skeleton microbiome communities. Based on their metabolic features, we characterize the carbon, sulfur, nitrogen cycling between *Halodesulfovibrio* and CAP, specifying the metabolic relationships among endolithic microbes in corals.

## Method

### Sample collection and anaerobic endolithic culturing

Three *Isopora palifera* colonies were collected from the ocean near Gonguuan (22°40’ N 121°27’ E) in Lyudao, Taiwan (also known as Green Island) on October 16, 2017. These colonies were placed in an anaerobic jar with an anaerobic pack immediately after sampling. Green layers from each colony were collected as described in our previous studies (11, 13). The anaerobic condition was maintained throughout the collection process. Bacteria in the green layers were enriched in the basal medium for *Prosthecochloris*, which consisted of 0.5 g/L KH_2_PO_4_, 5.3 g/L NaCl, 0.5 g/L MgSO_4_-7H_2_O, 0.7 g/L NH_4_Cl, 0.33 g/L KCl, 21 g/L Na_2_SO_4_, 4.0 g/L MgCl_2_-6H_2_O, 10 g/L NaHCO_3_, 0.07 g/L CaCl_2_-2H_2_O, and 0.005 g/L Resazurin, and supplemented with glucose (0.05%) as an additional carbon source (11, 40). The entire culturing process was performed under dim light (45.5 ± 31.5 lums/ft^2^) conditions.

### DNA extraction and whole-genome shotgun sequencing

Bacterial cells in the culture medium were centrifuged at 7,000 x *g* for 10 min at 20°C to obtain cell pellets. Total genomic DNA from the pellet was then extracted using the UltraClean Microbial DNA Isolation Kit (MioBio, Solana Beach, CA, USA) according to the manufacturer’s protocol and DNA concentration was determined by Nanodrop and Qubit. The DNA samples were sent to Yourgene Bioscience (Taipei, Taiwan) for library preparation and DNA sequencing by the Illumina MiSeq system (USA) with 2 x 300 cycles.

### Metagenome assembly and binning

Reads obtained from Illumina MiSeq were quality checked by FastQC (41). Quality trimming and removal of Illumina adaptors were performed by Trimmomatic v0.39 with following parameters: ILLUMINACLIP:TruSeq3-PE-2.fa:2:30:10:3: TRUE LEADING:10 TRAILING:10 SLIDINGWINDOW:5:15 MINLEN:50 CROP:300 (42). Leading and trailing bases with Phred quality score <15 were trimmed using a 5-base wide sliding window. Only reads with >50 bases were retained. The processed reads from three cultures were *de novo* assembled individually using megahit with k-mer sizes of 21, 31, 41, 51, 61, 71, 81, 91, and 99 (43) without scaffolding. Automated binning was performed using MetaBAT v0.32.5 with default settings, which reconstructed genomes from assembled metagenomic contigs based on probabilistic distances of genome abundance and tetranucleotide frequency (44).

### Quality assessment, taxonomic inference, and relative proportion of MAGs

The quality of each metagenome-assembled genome (MAG) was accessed by CheckM v1.0.13, which uses lineage-specific marker genes to estimate completeness and contamination (45). The taxonomy of each MAG was automatically assigned by GTDB-Tk v0.3.2 based on the placement of the genome in the reference tree, Average Nucleotide Identity (ANI) values, and relative evolutionary divergence (RED) values (46). To estimate the relative proportion of MAGs in each culture, reads were first mapped to assembled contigs using Bowtie2 v2.3.5 (47) with default settings. Mapped reads results were then used to obtain coverage for each contig and the relative proportion of each MAG with the ‘coverage’ and ‘profile’ command in CheckM, respectively.

### Genome annotation

The genome of coral-associated *Prosthecochloris* (CAP) and *Candidatus* Halodesulfovibrio lyudaonia were annotated using Prokka v1.13.7 with the ‘usegenus’ and ‘rfam’ options (48). The genomes were also annotated with KEGG functional orthologs (K numbers) by searching the putative protein sequences from Prokka against the KEGG database using BlastKoala (49). The K number annotation results were then used to reconstruct the transporter systems and metabolic pathways using KEGG mapper (50). Additionally, the transporter proteins were identified by searching for the putative protein sequences against TransportDB 2.0 (August 2019) using BLASTp (51).

### Recruitment of contigs with 16S rRNA gene sequences

The contigs with 16S rRNA gene sequences were originally not binned into the draft genome. To recruit the 16S rRNA gene, BLASTn was used to identify the contigs with *Prosthecochloris*-related 16S rRNA genes with an identity of >97%. Only one *Prosthecochloris*-related 16S rRNA gene was identified in each culture, consistent with the finding that only one CAP genome was recovered. Based on these results, the each contig containing *Prosthecochloris* 16S rRNA gene was moved into the CAP draft genomes.

### Average nucleotide identity (ANI) calculation and phylogenetic analysis

The ANIs between genomes were determined using the ANI calculator (52) and the ANI matrices were visualized using the pheatmap function (53) in R (R core team, 2016). To analyze the 16S rRNA gene phylogeny of *Chlorobiaceae* and *Halodesulfovibrio*, the available *Chlorobiaceae* genomes and representative *Desulfovibrio* genomes were retrieved from the RefSeq database (August 2019) (54) and 16S rRNA gene sequences in the genomes were extracted by Barrnap v0.9 (55). On the other hand, *Halodesulfovibrio* 16S rRNA gene sequences were downloaded from the NCBI 16S rRNA database and included in the analysis. A multiple sequence alignment of these 16S rRNA genes was performed using MUSCLE (56), followed by a tree reconstruction by the Maximum Likelihood method based on the Jukes-Cantor model and initial tree generation using the BioNJ method in MEGA7 (57, 58). The confidence levels of the tree were determined using 1000 bootstraps (59).

For the FMO phylogeny, the FMO proteins were retrieved from the available *Chlorobiaceae* genomes in RefSeq database (54). A tree was then inferred using the Maximum Likelihood method based on the JTT matrix-based model (60) and initial tree generation using the BioNJ method in MEGA7 (57) with 1000 bootstraps.

A tree was built from single-copy marker genes using the ezTree pipeline (61). Briefly, the putative genes in the genomes were identified by Prodigal (62), and the Pfam profiles of these genes were annotated using HMMER3 (63). Gene annotations were compared to identify single-copy marker genes among the input genomes. The amino acid sequences of single-copy marker genes were then aligned by MUSCLE (56). The alignments were trimmed using Gblocks (64), and a tree based on the concatenated alignment was constructed by Maximum Likelihood using FastTree with 1000 bootstraps (59, 65).

### Pan-genome analysis

Bacterial Pan Genome Analysis tool (BPGA) v1.3 (66) was used to perform a pan-genome analysis. The genes in the *Prosthecochloris* genomes were first clustered using USEARCH (67) with a 70% identity cutoff. Gene clusters present in all the genome were defined as core genes, and those present in at least two—but not all—of the genomes were defined as accessory genes. The representative sequences of CAP-specific accessory genes were then searched against the NCBI RefSeq database (54) to identify the potential orthologous genes in bacteria, with 40% identity and 50% alignment length cutoffs. In addition, the dN/dS values of each CAP-unique accessory gene were determined using the HyPhy tool in MEGA7 (57).

## Data Availability

Sequence reads of metagenomes have been submitted to NCBI sequence read archive (SRA) under SRA accession numbers SRR10714424, SRR10714423, SRR10714422, and SRR10714421, respectively. Supplementary data for this preprint is available on request to the corresponding author.

## Acknowledgements

This study was supported by funding from Academia Sinica. Y.H.C would like to acknowledge the Taiwan International Graduate Program (TIGP) for its fellowship towards his graduate studies. We would like to thank Noah Last of Third Draft Editing for his English language editing.

## Author contribution

Y.H.C, S.H.Y, and S.L.T conceived the idea for this study. Y.H.C and S.H.Y assembled the genomes, performed the bioinformatics analysis, and wrote the manuscript. K.T helped write the manuscript and modify the illustrations. C.Y.L and H.J.C collected coral skeleton samples and prepared the DNA samples. C.J.S provided the cultures. S.L.T supervised the overall study. All authors read and approved the manuscript.

## Supplemental Materials

**FIG S1. Heatmap of average nucleotide identity between two individual GSB genomes**.

Values of ANI < 70 are denoted as NA because values below 70% are not reliable.

**FIG S2. Pan-genome analysis**. (A) Core and Pan-Genome plot of *Prosthecochloris*. (B) Statistics from the pan-genome analysis, including number of core, unique, accessory, and exclusive absent genes.

**FIG S3**. COG (A) and KEGG (B) distributions of core, accessory, and unique genes from the pan-genome analysis.

**FIG S4. Phylogenetic tree based on core genome**. The protein sequences of 20 random orthologous gene clusters in the core genome were aligned by MUSCLE, and the tree were constructed and concatenated by the neighbor-joining method. The number of clade-specific accessory genes are shown in each branch.

**FIG S5. Molecular phylogenetic analysis of green sulfur bacteria**. (A) Phylogeny constructed from 16S rRNA from 20 *Desulfovibrio* genomes from the RefSeq database and *Halodesulfovibrio* using the maximum-likelihood method with 1000 bootstraps. 28 sequences and 1562 position were involved in the analysis. Scare bar represents 0.02 changes per nucleotide site. (B) Similarity matrix between each of the two SRB genomes created by Gegenees. The matrix was exported into a distance matrix, which was used to generate a dendrogram by SplitsTree 4 with the neighbor joining method. The scare bar indicates 1% difference among average BLASTN similarity scores.

**Table S1**. Mapped reads and inferred abundance of each bin in N1, N2, and N3 metagenomes.

**Table S2**. Genes present in CAP but absent in other *Chlorobi*.

**Table S3**. Metabolism pathways and ABC transporter in coral-associated *Prosthecochloris*.

**Table S4**. Metabolism pathways and transporter systems in *Halodesulfovibrio*.

**DATA SET S1**. Transporter genes annotated by transportDB 2.0 in coral-associated *Prosthecochloris* and *Ca*. H. lyudaonia.

